# SARS-CoV-2 Lambda variant exhibits higher infectivity and immune resistance

**DOI:** 10.1101/2021.07.28.454085

**Authors:** Izumi Kimura, Yusuke Kosugi, Jiaqi Wu, Daichi Yamasoba, Erika P Butlertanaka, Yuri L Tanaka, Yafei Liu, Kotaro Shirakawa, Yasuhiro Kazuma, Ryosuke Nomura, Yoshihito Horisawa, Kenzo Tokunaga, Akifumi Takaori-Kondo, Hisashi Arase, The Genotype to Phenotype Japan (G2P-Japan) Consortium, Akatsuki Saito, So Nakagawa, Kei Sato

**Affiliations:** Division of Systems Virology, Department of Infectious Disease Control, International Research Center for Infectious Diseases, The Institute of Medical Science, The University of Tokyo, Tokyo 1088639, Japan; Laboratory of Systems Virology, Institute for Frontier Life and Medical Sciences, Kyoto University, Kyoto 6068507, Japan; Graduate School of Pharmaceutical Sciences, Kyoto University, Kyoto 6068501, Japan; Department of Molecular Life Science, Tokai University School of Medicine, Kanagawa 2591193, Japan; CREST, Japan Science and Technology Agency, Saitama 3220012, Japan; Faculty of Medicine, Kobe University, Hyogo 6500017, Japan; Department of Veterinary Science, Faculty of Agriculture, University of Miyazaki, Miyazaki 8892192, Japan; Department of Immunochemistry, Research Institute for Microbial Diseases, Osaka University, Osaka 5650871, Japan; Laboratory of Immunochemistry, World Premier International Immunology Frontier Research Centre, Osaka University, Osaka 5650871, Japan; Department of Hematology and Oncology, Graduate School of Medicine, Kyoto University, Kyoto 6068507, Japan; Department of Pathology, National Institute of Infectious Diseases, Tokyo 1628640, Japan; Center for Infectious Disease Education and Research, Osaka University, Osaka 5650871, Japan; Center for Animal Disease Control, University of Miyazaki, Miyazaki 8892192, Japan; Graduate School of Medicine and Veterinary Medicine, University of Miyazaki, Miyazaki 8892192, Japan; Bioinformation and DDBJ Center, National Institute of Genetics, Mishima, Shizuoka 4118540, Japan

**Author notes:** Twitter: @SystemsVirology (Kei Sato). These authors contributed equally (Izumi Kimura, Yusuke Kosugi, Jiaqi Wu). Correspondences (S.N.), (K.Sato).

**Keywords:** SARS-CoV-2, COVID-19, spike protein, C.37, Lambda

## Abstract

SARS-CoV-2 Lambda, a new variant of interest, is now spreading in some South American countries; however, its virological features and evolutionary trait remain unknown. Here we reveal that the spike protein of the Lambda variant is more infectious and it is attributed to the T76I and L452Q mutations. The RSYLTPGD246-253N mutation, a unique 7-amino-acid deletion mutation in the N-terminal domain of the Lambda spike protein, is responsible for evasion from neutralizing antibodies. Since the Lambda variant has dominantly spread according to the increasing frequency of the isolates harboring the RSYLTPGD246-253N mutation, our data suggest that the insertion of the RSYLTPGD246-253N mutation is closely associated with the massive infection spread of the Lambda variant in South America.

**Highlights:**

- Lambda S is highly infectious and T76I and L452Q are responsible for this property
- Lambda S is more susceptible to an infection-enhancing antibody
- RSYLTPGD246-253N, L452Q and F490S confer resistance to antiviral immunity

**Graphical Abstract:** 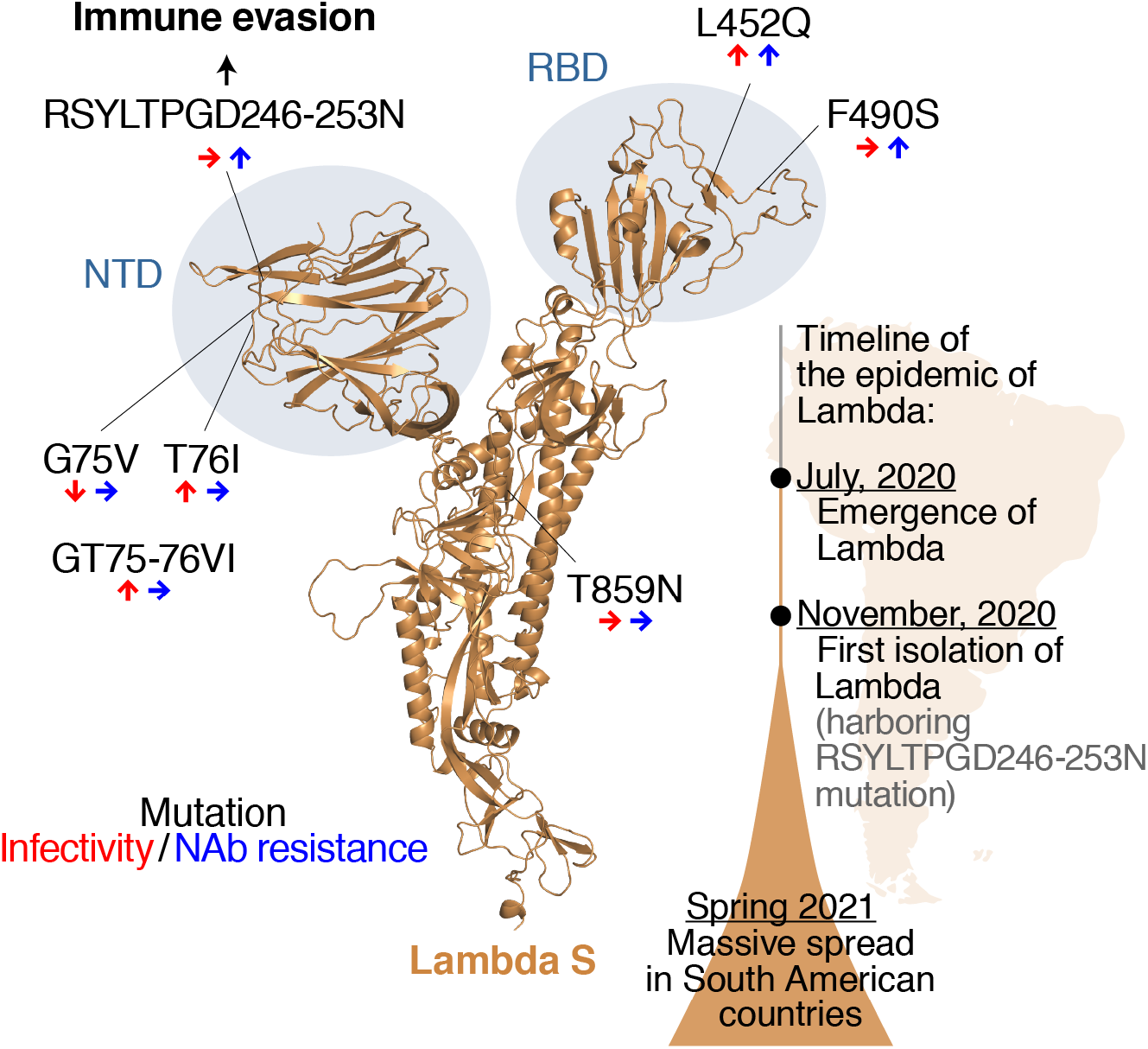

## Introduction

During the pandemic, severe acute respiratory syndrome coronavirus 2 (SARS-CoV-2), the causative agent of coronavirus disease 2019 (COVID-19), has been diversified. As of July 2021, there are four variants of concerns (VOCs), Alpha [B.1.1.7 lineage; the lineage classification is based on Phylogenetic Assignment of Named Global Outbreak (PANGO): https://cov-lineages.org/resources/pangolin.html], Beta (B.1.351 lineage), Gamma (P.1 lineage) and Delta (B.1.617.2 lineage), and four variants of interests (VOIs), Eta (B.1.525 lineage), Iota (B.1.526 lineage), Kappa (B.1.617.1 lineage) and Lambda (C.37 lineage) (WHO, 2021a). These variants are considered to be the potential threats to the human society.

VOCs and VOIs harbor multiple mutations in their spike (S) protein and are relatively resistant to the neutralizing antibodies (NAbs) that are elicited in convalescent and vaccinated individuals (Chen et al., 2021; Collier et al., 2021; Garcia-Beltran et al., 2021; Hoffmann et al., 2021; Liu et al., 2021a; Liu et al., 2021b; Planas et al., 2021; Wall et al., 2021a; Wang et al., 2021a; Wang et al., 2021b). Because the receptor binding domain (RBD) of the SARS-CoV-2 S protein is immunodominant, mutations in this domain can lead to the immune evasion (Piccoli et al., 2020). Additionally, the mutations in the N-terminal domain (NTD) of the SARS-CoV-2 S protein are associated with the escape neutralization (McCallum et al., 2021). Moreover, the antibodies that enhance viral infectivity [enhancing antibodies (EAbs)] were detected in severe COVID-19 patients, and these EAbs target NTD (Li et al., 2021; Liu et al., 2021c). Because natural mutations in the S NTD crucially influence the sensitivity to antibodies (Gobeil et al., 2021), the accumulation of mutations in this domain is closely associated with the infection spread of VOCs and VOIs.

The Lambda variant (also known as the C.37 lineage) is the newest VOI (designated on June 14, 2021) (WHO, 2021a) and is currently spreading in South American countries such as Peru, Chile, Argentina, and Ecuador (WHO, 2021a). Based on the information data from the Global Initiative on Sharing All Influenza Data (GISAID) database (https://www.gisaid.org; as of June 29, 2021), the Lambda variant has been isolated in 26 countries. Notably, the vaccination rate in Chile is relatively high; the percentage of the people who received at least one dose of COVID-19 vaccine was ~60% on June 1, 2021 (https://ourworldindata.org/covid-vaccinations). A recent paper also suggested that the vaccines have effectively prevented COVID-19 in Chile (Jara et al., 2021). Nevertheless, a big COVID-19 surge has occurred in Chile in Spring 2021 (WHO, 2021b), suggesting that the Lambda variant is proficient in escaping from the antiviral immunity elicited by vaccination. In this study, we reveal the evolutionary trait of the Lambda variant by molecular phylogenetic analysis. We further demonstrate that the RSYLTPGD246-253N mutation, a unique mutation in the NTD of the Lambda S protein, is responsible for the virological phenotype of the Lambda variant that can associate with the massive infection spread mainly in South American countries.

## Results

### Epidemic dynamics of the Lambda variant in South American countries

As of June 29, 2021, 1,908 genome sequences of the Lambda variant belonging to the PANGO C.37 lineage have been isolated from 26 countries and deposited in GISAID. Although it is considered that the Lambda variant was first detected in Peru in December 2020 (WHO, 2021a), our in-depth analysis revealed that the Lambda variant was first detected in Argentina on November 8, 2020 (GISAID ID: EPI_ISL_2158693) (**Figure 1A**; see **STAR★METHODS** for the detail). The fact that the percentage of the Lambda sequence is increasing in South American countries including Peru, Chile, and Argentina (**Figure 1A and Table S1**) suggests that the Lambda variant is spreading predominantly in these countries (**Table S2**). We then generated a maximum likelihood-based phylogenetic tree of the Lambda variant (C.37). Although there are some isolates that have been misclassified as the Lambda, which could be a sister group of the Lambda variant, our phylogenetic tree indicated the monophyly of genuine Lambda variant isolates (**Figure S1**).

**Figure 1.**
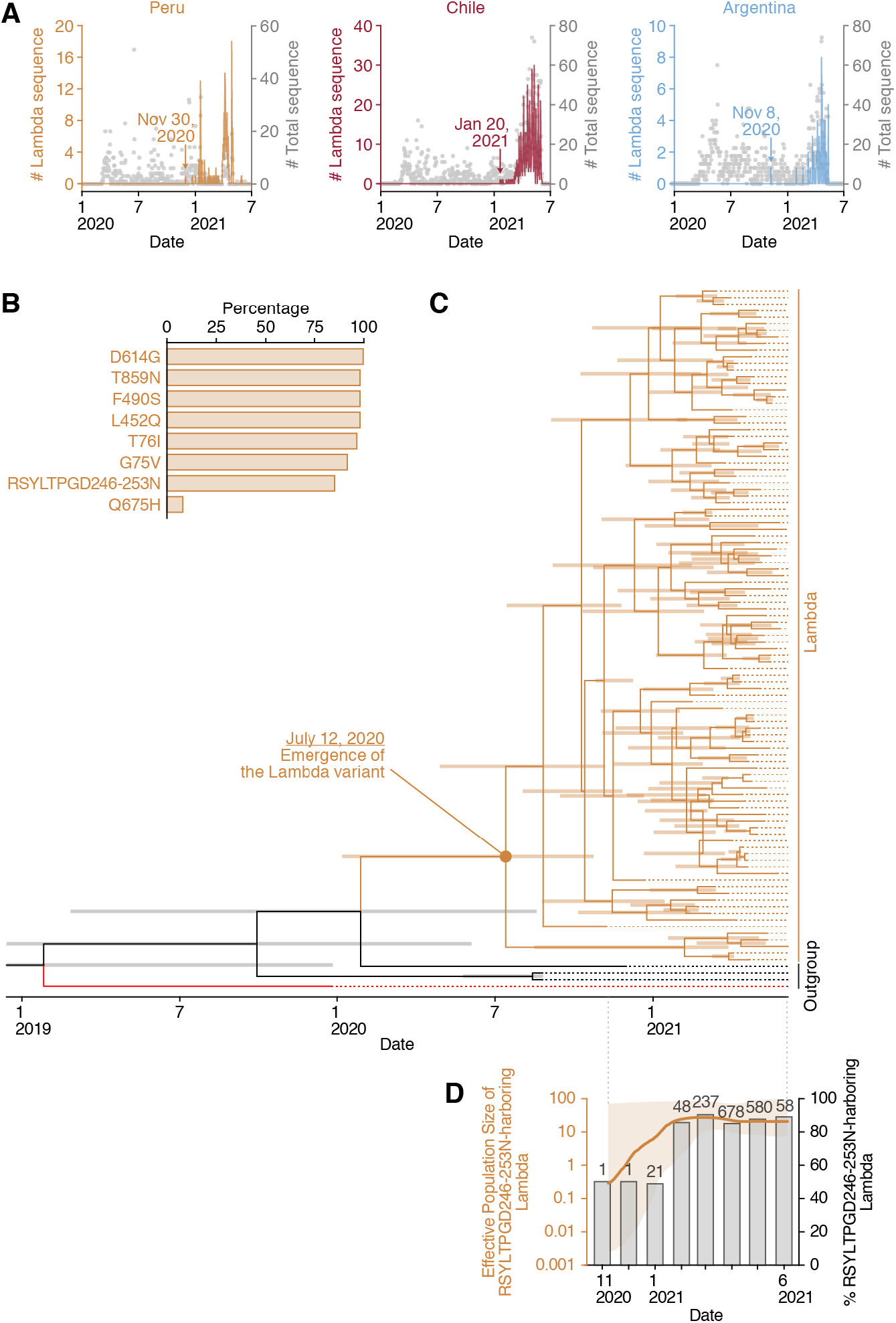
Epidemic and evolutionary dynamics of the Lambda variant. (**A**) Epidemic dynamics of the Lambda variant in three South American countries. The numbers of the Lambda variant (C.37 lineage) deposited per day from Peru (brown), Chile (dark red), and Argentina (pale blue) are indicated in lines. Grey dots indicate the numbers of SARS-CoV-2 genome sequences deposited in the GISAID database per day from respective countries. The raw data are summarized in **Tables S1 and S2**. (**B**) Proportion of amino acid replacements in the Lambda variant (C.37 lineage). The top 8 replacements conserved in the S protein of the Lambda variant (C.37 lineage) are summarized. The raw data are summarized in **Table S3**. (**C**) An evolutionary timetree of the Lambda variant (C.37 lineage). The estimated date of the emergence of the Lambda variant is indicated in the figure. The three sister sequences of the genuine C.37 lineage [GISAID ID: EPI_ISL_1532199 (B.1.1.1 lineage), EPI_ISL_1093172 (B.1.1.1 lineage) and EPI_ISL_1534656 (C.37 lineage)] are used as an outgroup and indicated in black. Wuhan-Hu-1 (GISAID ID: EPI_ISL_1532199), the oldest SARS-CoV-2 (isolated on December 26, 2019), is indicated in red. Bars on the internal nodes correspond to the 95% highest posterior density (HPD). The tree noted with the GISAID ID and sampling date at each terminal node is shown in **Figure S1B**. (**D**) Transition of the effective population size of the Lambda variant and the proportion of the Lambda variant harboring the RSYLTPGD246-253N mutation. The effective population size of the Lambda variant harboring the RSYLTPGD246-253N mutation (left y-axis) was analyzed by the Bayesian skyline plot. The initial date is when the first Lambda variant was sampled (November 8, 2020). The 95% highest posterior density (HPD) is shaded in brown. In the same panel, the proportion of the Lambda variants harboring RSYLTPGD246-253N mutation for each month (right y-axis) is also plotted. The number at each time point indicates the number of the Lambda variants harboring the RSYLTPGD246-253N mutation the mutation. The number in parentheses indicates the number of the Lambda variants deposited in the GISAID database. See also **Figure S1** and **Tables S1-S3**.

### Association of the lambda variant spread with the increasing frequency of the isolates harboring the RSYLTPGD246-253N mutation

The S protein of the consensus sequence of the Lambda variant bears six substitution mutations (G75V, T76I, L452Q, F490S, D614G and T859N) and a 7-amino-acid deletion in the NTD (RSYLTPGD246-253N) (**Table S3**). The analysis using the 1,908 sequences of the Lambda variant (C.37 lineage) showed that the six substitution mutations are relatively highly (> 90%) conserved (**Figure 1B** and **Table S3**). Although a large deletion in the NTD, the RSYLTPGD246-253N mutation, is also highly conserved, 287 out of the 1,908 sequences (15.0%) of the Lambda (C.37 lineage) genomes do not harbor this mutation (**Figure 1C** and **Table S3**). To ask whether the epidemic dynamics of the Lambda variant is associated with the emergence of the RSYLTPGD246-253N mutation, we examined all amino acid replacements in the S protein of the SARS-CoV-2 genomes deposited in the GISAID database (2,084,604 sequences; as of June 29, 2021). The RSYLTPGD246-253N mutation was first found in Argentina on November 8, 2020 (GISAID ID: EPI_ISL_2158693), which is the first isolate of the Lambda variant (**Figure 1A**), suggesting that this deletion event uniquely occurred in the ancestral lineage of the Lambda variant. We then analyzed the molecular evolutionary dynamics of the Lambda variants by performing the Bayesian tip-dating analysis. We showed that the Lambda variant bearing the RSYLTPGD246-253N mutation emerged around July 12, 2020 (95% CI, January 5, 2020 – October 22, 2020) (**Figure 1C** and **Figure S1B**). To infer the population dynamics of the lineage, we performed the Bayesian skyline plot analysis. This analysis showed that the effective population size of the Lambda variant has increased at the beginning of 2021 (**Figure 1D**). Intriguingly, when we plot the proportion of the Lambda variant that bears RSYLTPGD246-253N mutation, it was increased after the emergence of the Lambda variant and closely associated with the increase of effective population size (**Figure 1D**). These results suggest that the emergence of the RSYLTPGD246-253N mutation is associated with the outbreak of the Lambda variant in South America.

### Higher infectivity and resistance to NAbs of the Lambda variant

To address the virological phenotype of the Lambda variant, we prepared the viruses pseudotyped with the S proteins of the Lambda variant as well as four VOCs, Alpha (B.1.1.7), Beta (B.1.351), Gamma (P.1) and Delta (B.1.617.2) (**Table S4**). We also prepared the pseudoviruses with the S proteins of the D614G-bearing parental isolate (B.1) and the Epsilon (B.1.427/429) variant (**Table S4**), which is used to be a VOC/VOI by July 6, 2021 (WHO, 2021a), as controls of this experiment. As shown in **Figure 2A**, the infectivities of the Alpha and Beta variants were significantly lower than that of the parental D614G S, and the infectivities of Gamma variant and the parental D614G S were comparable. On the other hand, the infectivities of the Delta, Epsilon and Lambda variants were significantly higher than that of the parental D614G (**Figure 1A**). This pattern was independent of the input dose of pseudoviruses used (**Figure S2A**).

**Figure 2.**
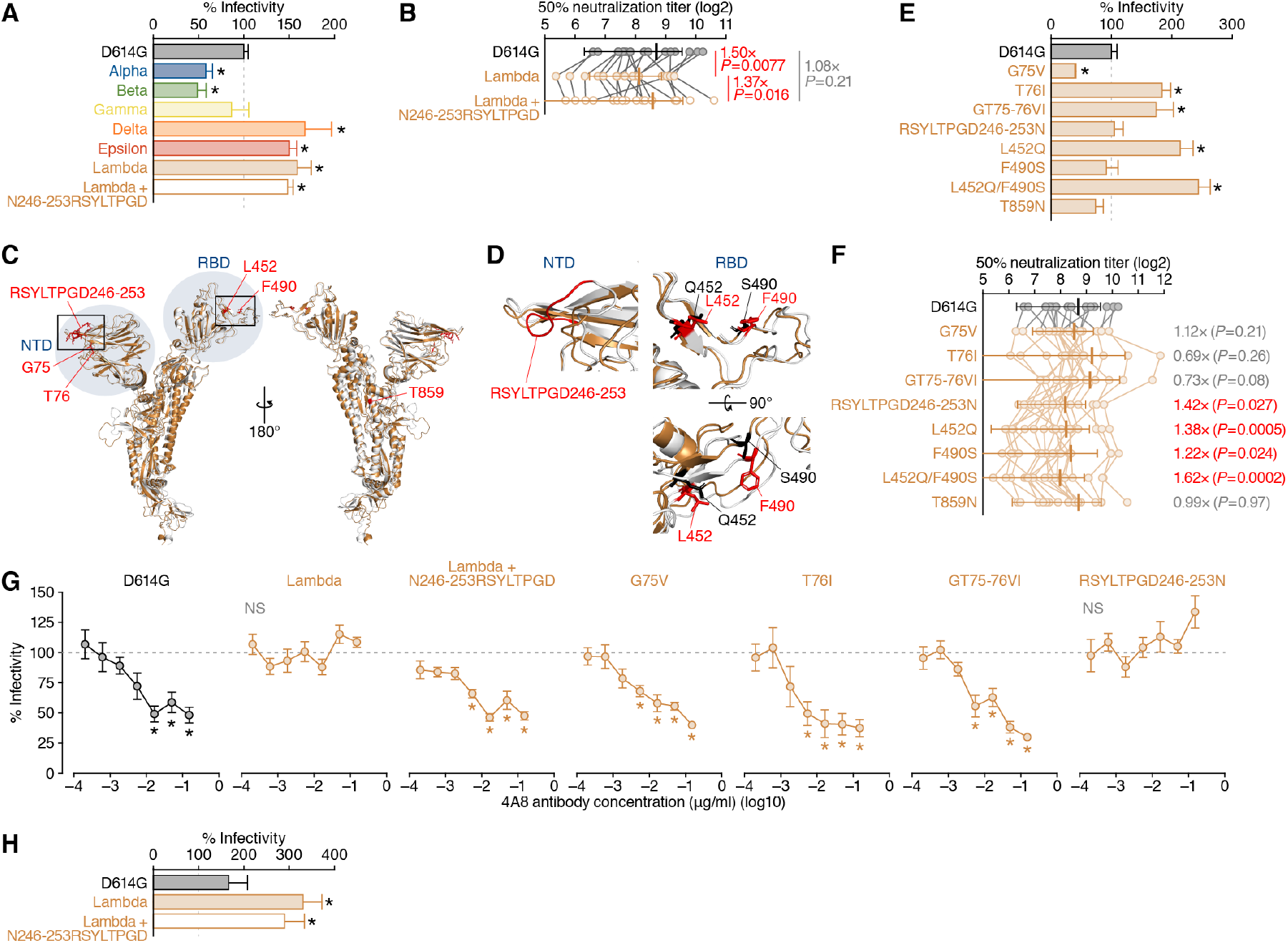
Virological and immunological features of the Lambda variant. (**A**) Pseudovirus assay. The HIV-1-based reporter viruses pseudotyped with the SARS-CoV-2 S proteins of the parental D614G (B.1), Alpha (B.1.1.7), Beta (B.1.351), Gamma (P.1), Delta (B.1.617.2), Epsilon (B.1.427), Lambda (C.37) variants as well as the Lambda+N246-253RSYLTPGD derivative were prepared as described in **STAR★METHODS**. The mutations in each variant are listed in **Table S4**. The pseudoviruses were inoculated into HOS-ACE2/TMPRSS2 cells at 1,000 ng HIV-1 p24 antigen, and the percentages of infectivity compared to the virus pseudotyped with parental S D614G are shown. (**B**) Neutralization assay. Neutralization assay was performed using the pseudoviruses with the S proteins of parental D614G, Lambda and Lambda+N246-253RSYLTPGD and 18 BNT162b2-vaccinated sera as described in **STAR★METHODS**. The raw data are shown in **Figure S2B**. The number in the panel indicates the fold change of neutralization resistance of the Lambda S to the D614G S or the Lambda+N246-253RSYLTPGD derivative. (**C and D**) Structural insights of the mutations in the Lambda S. (**C**) Overlaid overviews of the crystal structure of SARS-CoV-2 S (PDB: 6ZGE, white) (Wrobel et al., 2020) and a homology model of the Lambda S (brown). The mutated residues in the Lambda S and the regions of NTD and RBD are indicated in red and blue. The squared regions are enlarged in (**D**). Mutated residues in the NTD (left) and RBD (right) of the Lambda S. The residues in the parental S and the Lambda S are indicated in red and black. (**E**) Pseudovirus assay. The HIV-1-based reporter viruses pseudotyped with the SARS-CoV-2 S proteins bearing respective mutations of the Lambda variant as well as the D614G S were prepared. The pseudoviruses were inoculated into HOS-ACE2/TMPRSS2 cells at 1,000 ng HIV-1 p24 antigen, and the percentages of infectivity compared to the virus pseudotyped with parental S D614G are shown. (**F**) Neutralization assay. Neutralization assay was performed using the pseudoviruses used in **Figure 2B** and 18 BNT162b2-vaccinated sera as described in **STAR★METHODS**. The raw data are shown in **Figure S2B**. The number in the panel indicates the fold change of neutralization resistance to the D614G S. (**G and H**) Effect of monoclonal antibodies. (**G**) Antiviral effect of an NTD-targeting NAb clone 4A8 (Chi et al., 2020). (**H**) Enhancing effect of an EAb clone COV2-2490 (Liu et al., 2021c). The percentages of infectivity compared to the virus without antibodies are shown. In **A, E and H**, assays were performed in triplicate, and the average is shown with SD. Statistically significant differences (*, *P* < 0.05) versus the D614G S were determined by Student’s *t* test. In **B and F**, assays were performed in triplicate, and the average is shown with SD. Statistically significant differences were determined by Wilcoxon matched-pairs signed rank test. The *P* values are indicated in the figure. In **G**, assays were performed in quadruplicate, and the average is shown with SD. Statistically significant differences (*, *P* < 0.05) versus the value without antibody were determined by Student’s *t* test. NS, no statistical significance. See also **Figure S2 and Table S4**.

To assess the effect of a characteristic mutation of the Lambda variant, the RSYLTPGD246-253N mutation (**Figure 1**), on viral infectivity, we prepared the pseudovirus with the Lambda S derivative recovering this deletion mutation (“Lambda+N246-253RSYLTPGD”). The infectivity of this mutant was comparable to that of the Lambda S (**Figures 2A and S2A**), suggesting that the 7-amino-acid deletion in the NTD does not affect viral infection.

Because the NAb resistance is a remarkable phenotype of most VOCs [reviewed in (Harvey et al., 2021)], we next analyzed the sensitivity of the Lambda S to the NAbs induced by BNT162b2 vaccination. As shown in **Figure 2B**, the Lambda S is 1.5-fold in average (2.63-fold at a maximum) more resistant to the BNT162b2-induced antisera than the parental D614G S (*P* = 0.0077 by Wilcoxon matched-pairs signed rank test). On the other hand, the neutralization level of the Lambda+N246-253RSYLTPGD was similar to the D614G pseudovirus (*P* = 0.21 by Wilcoxon matched-pairs signed rank test), and this recovered variant was 1.37-fold more sensitive to the vaccine-induced neutralization than the Lambda (*P* = 0.0016 by Wilcoxon matched-pairs signed rank test) (**Figure 2B**). These results suggest that the Lambda S is highly infectious and resistant to the vaccine-induced humoral immunity, and the robust resistance of the Lambda S to the vaccine-induced neutralization is determined by a large deletion in the NTD.

### Effect of the consensus mutations in the Lambda S on viral infectivity and NAb sensitivity

We next plotted six substitution mutations (G75V, T76I, L452Q, F490S, D614G and T859N) and a deletion mutation (RSYLTPGD246-253N) of the Lambda variant on the structure of the SARS-CoV S protein. Structural analysis showed that the three mutations, G75V, T76I and RSYLTPGD246-253N are in the NTD (**Figure 2C**), and the RSYLTPGD246-253N mutation is located in a loop structure, which is designated as loop 5 structure (residues 246-260) in a previous study (Chi et al., 2020) (**Figure 2D**). The L452Q and F490S mutations are in the RBD (**Figures 2C and 2D**), but neither residues are located on the ACE2 interface (**Figure S2D**). The T859N mutation is in the heptad repeat 1 of the S2 subunit (**Figure 2C**).

To investigate the effects of these seven consensus mutations in the Lambda S on viral infectivity and NAb sensitivity, we prepared the viruses pseudotyped with the D614G S-based derivatives that possess respective mutations of the Lambda variant. **Figure 2E** showed that the G75V mutation significantly reduces viral infectivity, while the T76I and GT75-76VI mutations significantly increases it. Additionally, 91.5% (1,746/1,908) of the Lambda variant sequence possesses these two mutations, and the phylogenetic tree of the Lambda variant indicated that the variant harboring either G75V or T76I sporadically emerged during the epidemic of the Lambda variant (**Figure S1A**). These findings suggest that T76I is a compensatory mutation to recover the decreased infectivity by the G75V mutation.

Similar to the experiment using the pseudoviruses with the Lambda S and Lambda+N246-253RSYLTPGD (**Figure 2A**), the insertion of the RSYLTPGD246-253N mutation did not affect viral infectivity (**Figure 2E**). The infectivity of the T859N mutation was also similar to that of the parental D614G pseudovirus (**Figure 2E**). When we focus on the effect of the mutations in the RBD, the L452Q and L452Q/F490S mutations significantly increased viral infectivity, while the F490S sole mutation did not (**Figure 2E**). Taken together, these results suggest that the T76I and L452Q mutations are responsible for the higher infectivity of the Lambda S (**Figure 2A**). The effect of each mutation was similar in different amounts of pseudoviruses used and in the target cells without TMPRSS2 expression (**Figure S2E**).

We next assessed the sensitivity of these pseudoviruses with the mutated S proteins to BNT162b2-induced antisera. As shown in **Figure 2F**, the G75V, T76I, GT75-76VI and T852N mutations did not affect the vaccine-induced neuralization. On the other hand, the RSYLTPGD246-253N mutant exhibited a significant resistance to the vaccine-induced neuralization (*P* = 0.027 by Wilcoxon matched-pairs signed rank test; **Figure 2F**), which is relevant to the experiment with the S proteins of the Lambda variant and the Lambda+N246-253RSYLTPGD derivative (**Figure 2B**). Additionally, we found that the L452Q and F490S mutations confer resistance to the vaccine-induced antisera (**Figure 2F**). The results that the F490S mutation does not affect viral infectivity (**Figure 2E**) but confers the resistance to the vaccine-induced antisera (**Figure 2F**) suggest that this mutation has acquired to be resistant to antiviral humoral immunity. On the other hand, the L452Q mutation not only increases viral infectivity (**Figure 2E**) but also augments the resistance to the vaccine-induced antisera (**Figure 2F**), suggesting that that this mutation can be critical for the viral dissemination in the human population.

To further assess the association of the mutations in the Lambda S, particularly those in the NTD, we used two monoclonal antibodies that targets the NTD: an NTD-targeting NAb, clone 4A8 (Chi et al., 2020), and an EAb, clone COV2-2490, that recognizes the NTD and enhances viral infectivity (Liu et al., 2021c). As shown in **Figure 2G**, the 4A8 antibody inhibited the pseudovirus infections of the parental D614G, Lambda+N246-253RSYLTPGD derivative, G75V, T76I and GT75-75VI in dose-dependent manners. Intriguingly, the pseudovirues with the S proteins of the Lambda and the RSYLTPGD246-253N mutant were resistant to the antiviral effect mediated by the 4A8 antibody (**Figure 2G**). These results suggest that the RSYLTPGD246-253N mutation critically affects the sensitivity to certain NAbs targeting the NTD. On the other hand, the COV2-2490 antibody enhanced the infectivities of the parental D614G, the Lambda, and the Lambda+N246-253RSYLTPGD derivative (**Figure 2H**). Particularly, the infectivities of the Lambda and the Lambda+N246-253RSYLTPGD derivative were more significantly enhanced than the parental D614G (**Figure 2H**). These data suggest that the Lambda S is more susceptible to the EAb-mediated virus infection enhancement.

## Discussion

In this study, we demonstrated that three mutations, the RSYLTPGD246-253N, L452Q and F490S mutations, respectively confer resistance to the vaccine-induced antiviral immunity. Additionally, the T76I and L452Q mutations contribute to enhanced viral infectivity. Our data suggest that there are at least two virological features on the Lambda variant: increasing viral infectivity (by the T76I and L452Q mutations) and exhibiting resistance to antiviral immunity (by the RSYLTPGD246-253N, L452Q and F490S mutations).

Virological experiments demonstrated that a large 7-amino-acid deletion, the RSYLTPGD246-253N mutation, does not affect viral infectivity but is responsible for the resistance to the vaccine-induced neutralization as well as an NTD-targeting NAb. Additionally, molecular phylogenetic analyses showed that the transition of the proportion of the Lambda variant harboring a large 7-amino-acid deletion, the RSYLTPGD246-253N mutation, is associated with the increase of the effective population size of this variant. Therefore, the emergence of the RSYLTPGD246-253N mutation could be one of a driving forces behind the spread of this variant in the human population. In fact, here we showed that the Lambda S is more resistant to the vaccine-induced antisera than the Lambda+N246-253RSYLTPGD S derivative. Our results suggest that the resistance of the Lambda variant against antiviral humoral immunity was conferred by the RSYLTPGD246-253N mutation. Importantly, the RSYLTPGD246-253N mutation overlaps with a component of the NTD “supersite” (Chi et al., 2020). Altogether, these observations suggest that the NTD “supersite” is immunodominant and closely associate with the efficacy of the vaccine-induced neutralization, and further support the possibility that the emergence of the RSYLTPGD246-253N mutation triggered the massive spread of the Lambda variant.

We showed that the infectivity of the viruses pseudotyped with the Lambda S is significantly higher than that with the parental D614G S. Additionally, consistent with our previous reports (Mlcochova et al., 2021; Motozono et al., 2021b), the infectivity of the viruses pseudotyped with the S proteins of the Delta and Epsilon variants was significantly higher than that of the parental D614G S. A common feature of the Lambda, Delta and Epsilon variants is the substitution in the L452 of SARS-CoV-2 S protein: the Lambda variant harbors the L452Q mutation, while the Delta and Epsilon variants possess the L452R mutation (Mlcochova et al., 2021; Motozono et al., 2021b). Here we demonstrated that the L452Q mutation increases viral infectivity. Together with our previous observation that the L452R mutation enhances viral infectivity (Motozono et al., 2021b), it is strongly suggested that the relatively higher infectivity of the Lambda, Delta and Epsilon variants is attributed to the L452Q/R mutation. The fact that the Lambda and Delta variants are currently a VOI and a VOC, respectively, suggests that their increasing spread in the human population is partly attributed to their higher infectivity compared to the parental SARS-CoV-2. In contrast, nevertheless of its higher infectivity, the Epsilon variant, an ex VOC, has been excluded from the VOC/VOI classification on July 6, 2021 because this variant was stamped out (WHO, 2021a). The transient and unsuccessful (compared to the other VOCs) spread of the Epsilon variant in the human population implies that increasing viral infectivity is insufficient to maintain efficient spread in the human population. In addition to increasing viral infectivity, the Delta variant exhibits higher resistance to the vaccine-induced neutralization (Mlcochova et al., 2021; Wall et al., 2021b). Similarly, here we showed that the Lambda variant equips not only increased infectivity but also resistance against antiviral immunity. These observations suggest that acquiring at least two virological features, increased viral infectivity and evasion from antiviral immunity, is pivotal to the efficient spread and transmission in the human population.

Gobeil et al. have suggested that the mutations in the S NTD drive viral transmission and escape from antiviral immunity (Gobeil et al., 2021). In fact, three out of the current four VOCs harbor the deletions in the S NTD [reviewed in (Harvey et al., 2021)]. The Alpha variant bears the 2-amino-acid deletion, the HV69-70 deletion, in the NTD. Meng et al. showed that the HV69-70 deletion does not affect the NAb sensitivity but increase viral infectivity (Meng et al., 2021), suggesting that the virological significance of the deletion of a portion of the NTD between the Alpha (the HV69-70 deletion) and the Lambda (the RSYLTPGD246-253N mutation) is different. A cluster of the Beta variant also possesses the 3-amino-acid deletion, the LAL242-244 deletion. This deletion does not critically affect the sensitivity to the vaccine-induced neutralization but exhibits resistance to some NTD-targeting NAbs such as 4A8 (Liu et al., 2021b; Wang et al., 2021b). Interestingly, the Delta variant (B.1.617.2 lineage), an VOC, harbors the 2-amino-acid deletion, the EFR156-8G mutation, while its relative VOI, the Kappa variant (B.1.617.1 lineage), does not (WHO, 2021a). Although the virological significance of the EFR156-8G mutation remains unclear, this mutation in the S NTD may associate with the spread of the Delta variant worldwide. Together with our findings that the RSYLTPGD246-253N confers resistance to the vaccine-induced antisera and an NTD-targeting NAb, 4A8, the accumulative mutations in the S NTD may closely associate with the virological feature of the variants that can explain their spreading efficacy in the human population. Particularly, 4A8 targets the NTD “supersite” (Harvey et al., 2021; Lok, 2021) that includes RSYLTPGD246-253 (Chi et al., 2020). Because the RSYLTPGD246-253N mutation is responsible for the resistance to the vaccine-induced antiviral effect of the Lambda variant, our data support the possibility that the NTD “supersite” is immunodominant and acquiring immune escape mutations in this region associates with the efficacy of viral dissemination in the human population, which is proposed in previous reports (Harvey et al., 2021; McCallum et al., 2021). Moreover, we revealed that at least an EAb, COV2-2490, more preferentially enhances the Lambda S-mediated infection. Although the enhancing effect is independent of the RSYLTPGD246-253N mutation, such enhancement may associate with feasible spread of the Lambda variant.

By molecular phylogenetic analyses and virological experiments, here we elucidated how the Lambda variant was originated and acquired virological properties. Because the Lambda variant is a VOI, it might be considered that this variant is not an ongoing threat compared to the pandemic VOCs. However, because the Lambda variant is relatively resistant to the vaccine-induced antisera, it might be possible that this variant is feasible to cause breakthrough infection (Hacisuleyman et al., 2021; Jacobson et al., 2021; Nixon and Ndhlovu, 2021; Rana et al., 2021). Moreover, elucidating the evolutionary trait of threatening SARS-CoV-2 variants can explain the possibility to lead to wider epidemic, and revealing the virological features of the respective mutations acquired in VOCs and VOIs should be important to prepare the risk of newly emerging SARS-CoV-2 variants in the future.

## STAR★METHODS

- KEY RESOURCES TABLE
- RESOURCE AVAILABILITY
  ○ Lead Contact
  ○ Materials Availability
  ○ Data and Code Availability
- EXPERIMENTAL MODEL AND SUBJECT DETAILS
  ○ Ethics Statement
  ○ Collection of BNT162b2-Vaccinated Sera
  ○ Cell Culture
- METHOD DETAILS
  ○ Viral Genome Sequence Analysis
  ○ Plasmid Construction
  ○ Protein Structure Homology Model
  ○ Pseudovirus Assay
  ○ Antibody Treatment
- QUANTIFICATION AND STATISTICAL ANALYSIS

## Supplemental Information

Supplemental Information includes 2 figures and 5 tables and can be found with this article online.

## Consortia

The Genotype to Phenotype Japan (G2P-Japan) Consortium: Mika Chiba, Shigeru Fujita, Hirotake Furihata, Naoko Misawa, Nanami Morizako, Akiko Oide, Mai Suganami, Miyoko Takahashi, Miyabishara Yokoyama

## Acknowledgments

We would like to thank all members belonging to The Genotype to Phenotype Japan (G2P-Japan) Consortium. The super-computing resource was provided by Human Genome Center at The University of Tokyo and the NIG supercomputer at ROIS National Institute of Genetics.

This study was supported in part by AMED Research Program on Emerging and Re-emerging Infectious Diseases 20fk0108163 (to A.S.), 20fk0108146 (to K.Sato), 20fk0108270 (to K.Sato) and 20fk0108413 (to S.N. and K.Sato); AMED Research Program on HIV/AIDS 21fk0410033 (to A.S.) and 21fk0410039 (to K.Sato); AMED Japan Program for Infectious Diseases Research and Infrastructure 20wm0325009 and 21wm0325009 (to A.S.); JST SICORP (e-ASIA) JPMJSC20U1 (to K.Sato); JST SICORP JPMJSC21U5 (to K.Sato), JST CREST JPMJCR20H6 (to S.N.) and JPMJCR20H4 (to K.Sato); JSPS KAKENHI Grant-in-Aid for Scientific Research C 19K06382 (to A.S.), Scientific Research B 18H02662 (to K.Sato) and 21H02737 (to K.Sato); JSPS Fund for the Promotion of Joint International Research (Fostering Joint International Research) 18KK0447 (to K.Sato); JSPS Core-to-Core Program JPJSCCA20190008 (A. Advanced Research Networks) (to K.Sato); JSPS Research Fellow DC1 19J20488 (to I.K.); ONO Medical Research Foundation (to K.Sato); Ichiro Kanehara Foundation (to K.Sato); Lotte Foundation (to K.Sato); Mochida Memorial Foundation for Medical and Pharmaceutical Research (to K.Sato); Daiichi Sankyo Foundation of Life Science (to K.Sato); Sumitomo Foundation (to K.Sato); Uehara Foundation (to K.Sato); Takeda Science Foundation (to K.Sato); The Tokyo Biochemical Research Foundation (to K.Sato); a Grant for Joint Research Projects of the Research Institute for Microbial Diseases, Osaka University (to A.S.); and Joint Usage/Research Center program of Institute for Frontier Life and Medical Sciences, Kyoto University (to K.Sato).

---

**Figure S1.**
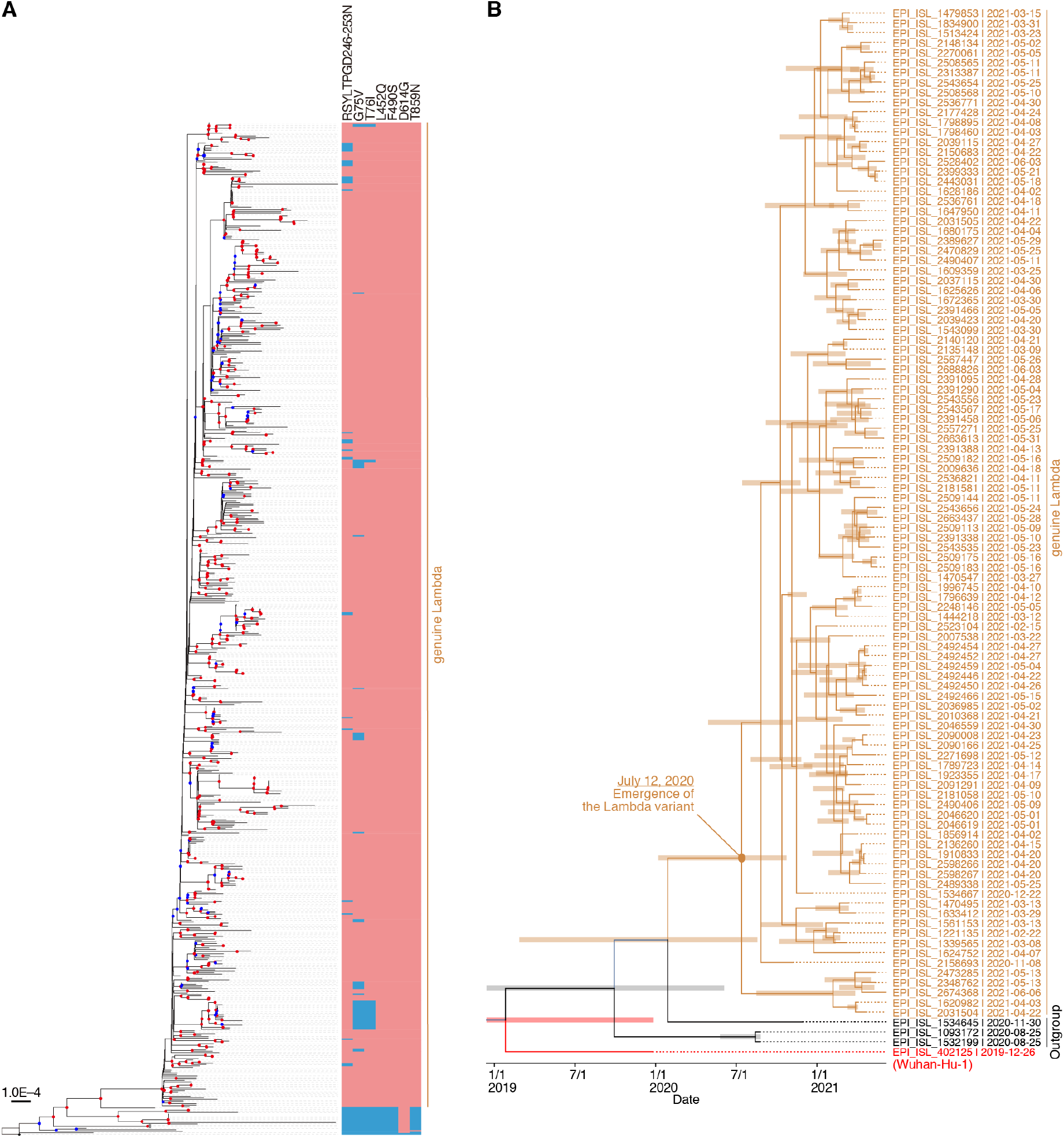
A maximum likelihood-based phylogenetic tree and an evolutionary timetree of the Lambda variant (C.37 lineage) (Related to Figure 1). (**A**) The 696 SARS-CoV-2 genome sequences were used for the analysis. Wuhan-Hu-1 (GISAID ID: EPI_ISL_1532199), located in the bottom of the tree, was indicated by a black star. EPI_ISL_1532199 and EPI_ISL_1093172 belonging to the B.1.1.1 lineage were indicated by grey stars. Red or blue circle on the branch was shown in each internal node if the bootstrap value was ≥ 80 or ≥ 50 (n = 1,000). A color box in pink or pale blue indicates the mutation in the S protein exist or not, respectively. (**B**) An evolutionary timetree of the Lambda variant (C.37 lineage). The estimated date of the emergence of the Lambda variant is indicated in the figure. The GISAID ID and sampling date is noted at each terminal node. The three sister sequences of the genuine C.37 lineage [GISAID ID: EPI_ISL_1532199 (B.1.1.1 lineage), EPI_ISL_1093172 (B.1.1.1 lineage) and EPI_ISL_1534656 (C.37 lineage)] are used as an outgroup and indicated in black. Wuhan-Hu-1 (GISAID ID: EPI_ISL_1532199), the oldest SARS-CoV-2 (isolated on December 26, 2019), is indicated in red. Bars on the internal nodes correspond to the 95% highest posterior density (HPD).

**Figure S2.**
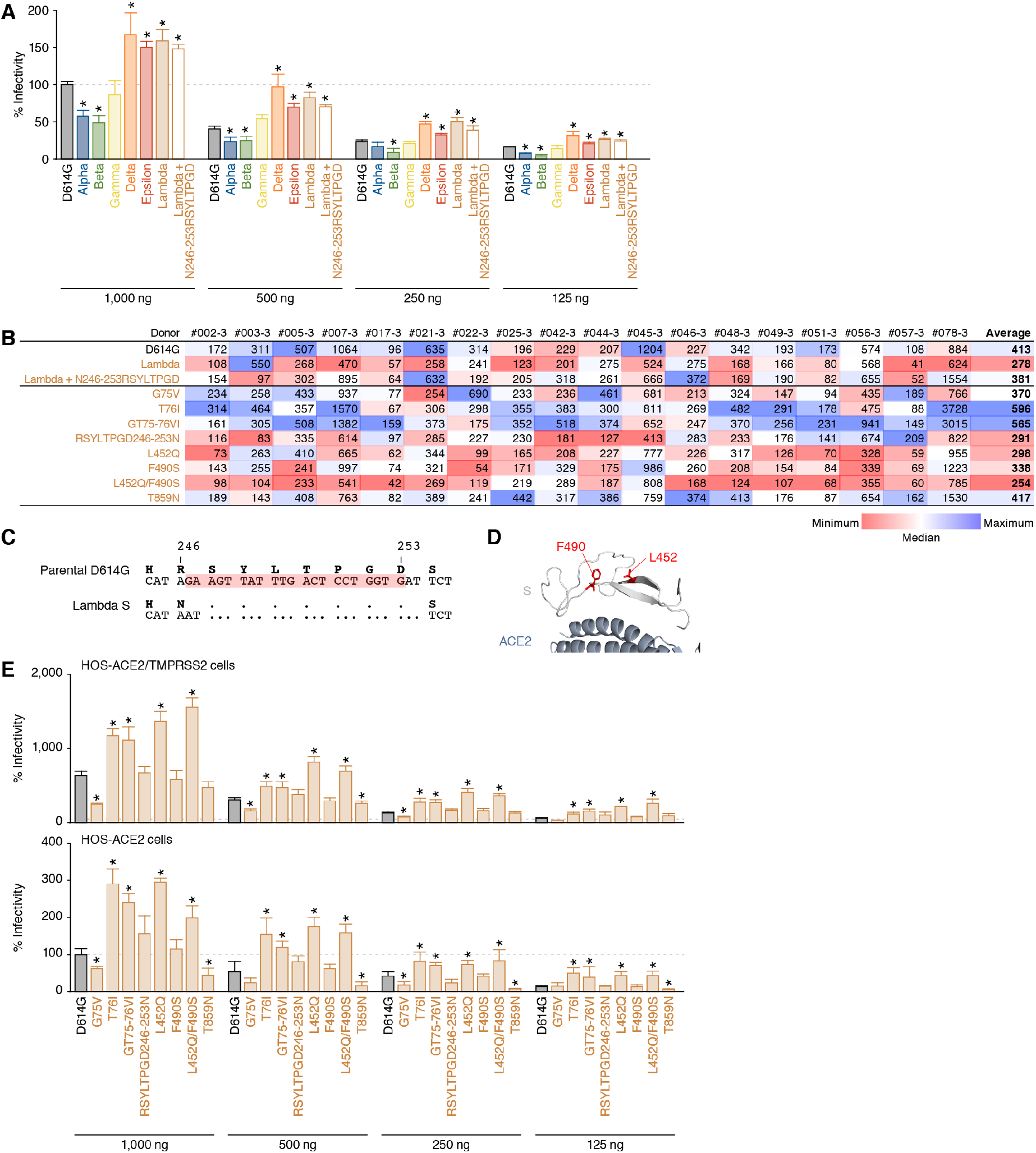
Virological features of the Lambda S (Related to Figure 2). (**A**) Pseudovirus assay. The HIV-1-based reporter viruses pseudotyped with the SARS-CoV-2 S proteins of the parental D614G (B.1), Alpha (B.1.1.7), Beta (B.1.351), Gamma (P.1), Delta (B.1.617.2), Epsilon (B.1.427) and Lambda (C.37) variants as well as the Lambda+N246-253RSYLTPGD derivative were prepared as described in **STAR★METHODS**. The pseudoviruses were inoculated into HOS-ACE2/TMPRSS2 cells at 4 different doses (1,000, 500, 250 or 125 ng HIV-1 p24 antigen), and percentages of infectivity compared to the virus pseudotyped with parental S D614G (1,000 ng HIV-1 p24 antigen) are shown. Assays were performed in triplicate. Note that the data of 1,000 ng HIV-1 p24 antigen are identical to those shown in **Figure 1B**. In **A**, assays were performed in triplicate, and the average is shown with SD. Statistically significant differences (*, *P* < 0.05) versus the D614G S were determined by Student’s *t* test. (**B**) Neutralization assay. Eighteen vaccinated sera were used for the neutralization assay. The 50% neutralization titers of respective serum against respective virus are shown. The values are summarized in **Figures 2B and 2F**. (**C**) Amino acid and nucleotide sequences of the residues 245-254 of the SARS-CoV-2 S. The amino acid sequences (residues 245-254, bold) and nucleotide sequences of the parental S (top) and the Lambda S (bottom) are shown. The nucleotides shaded in red in the parental S are deleted in the Lambda S, resulting in the RSYLTPGD246-253N mutation. (**D**) Positions of the residues L452 and F490. The residues L452Q and F490S are labeled in the cocrystal structure of SARS-CoV-2 and human ACE2 (PDB: 6M17) (Yan et al., 2020) in red. (**E**) Pseudovirus assay. The HIV-1-based reporter viruses pseudotyped with the SARS-CoV-2 S proteins bearing respective mutations of the Lambda variant as well as the D614G S were prepared. The pseudoviruses were inoculated into HOS-ACE2/TMPRSS2 cells (top) or HOS-ACE2 cells (bottom) at 4 different doses (1,000, 500, 250 or 125 ng HIV-1 p24 antigen), and percentages of infectivity compared to the virus pseudotyped with parental S D614G (1,000 ng HIV-1 p24 antigen) in HOS-ACE2 cells are shown. Assays were performed in triplicate. Note that the data of 1,000 ng HIV-1 p24 antigen in HOS-ACE2/TMPRSS2 cells are identical to those shown in **Figure 2E**. In **E**, statistically significant differences (*, *P* < 0.05) versus the D614G S were determined by Student’s *t* test.

**Table S1.** Frequency of the Lambda variant (C.37 lineage) and all SARS-CoV-2 genomes sampled in Chile, Argentina, and Peru per day, related to Figure 1

**Table S2.** Number of the Lambda variant sequences deposited from 26 countries, related to Figure 1

**Table S3.** Mutations in the S proteins of the Lambda variants obtained from the GISAID database (as of June 29, 2021), related to Figure 1

**Table S4.** Mutations in the S proteins of SARS-CoV-2 variants used in this study, related to Figure 2

**Table S5.** Primers for the construction of S derivatives, related to Figure 2

## STAR★METHODS

### KEY RESOURCES TABLE

### RESOURCE AVAILABILITY

#### Lead Contact

Further information and requests for resources and reagents should be directed to and will be fulfilled by the Lead Contact, Kei Sato.

#### Materials Availability

All unique reagents generated in this study are listed in the Key Resources Table and available from the Lead Contact with a completed Materials Transfer Agreement.

#### Data and Code Availability

Additional Supplemental Items are available from Mendeley Data at http://...

### EXPERIMENTAL MODEL AND SUBJECT DETAILS

#### Ethics Statement

For the use of human specimen, all protocols involving human subjects recruited at Kyoto University were reviewed and approved by the Institutional Review Boards of Kyoto University (approval number G0697). All human subjects provided written informed consent.

#### Collection of BNT162b2-Vaccinated Sera

Peripheral blood were collected four weeks after the second vaccination of BNT162b2 (Pfizer-BioNTech), and the sera of 18 vaccinees (average age: 40, range: 28-59, 22% male) were isolated from peripheral blood. Sera were inactivated at 56°C for 30 min and stored at –80°C until use.

#### Cell Culture

HEK293T cells (a human embryonic kidney cell line; ATCC CRL-3216), and HOS cells (a human osteosarcoma cell line; ATCC CRL-1543) were maintained in Dulbecco’s modified Eagle’s medium (high glucose) (Wako, Cat# 044-29765) containing 10% fetal calf serum and 1% PS.

HOS-ACE2/TMPRSS2 cells, the HOS cells stably expressing human ACE2 and TMPRSS2, were prepared as described previously (Ferreira et al., 2021; Ozono et al., 2021).

HOS-ACE2 cells, the HOS cells stably expressing human ACE2, were prepared as described previously (Saito et al., 2021).

### METHOD DETAILS

#### Viral Genome Sequence Analysis

All SARS-CoV-2 genome sequences and annotation information used in this study were downloaded from GISAID (https://www.gisaid.org) as of June 29, 2021 (2,084,604 sequences). We obtained 1,908 genomes of SARS-CoV-2 Lambda variant (C.37 lineage based on the PANGO annotation) in the GISAID metadata. We confirmed all of them are isolated from humans. To estimate when a Lambda variant harboring the RSYLTPGD246-253N deletion mutation in the S protein occurred, we screened 1,908 Lambda variants by removing genomes 1) containing more than 5 undetermined nucleotides at coding regions and 2) having an unknown sampling date. We then collected 644 and 49 viral genomes with and without RSYLTPGD246-253N deletion mutation in the Lambda S protein. We used Wuhan-Hu-1 strain isolated in China on December 31, 2019 (GenBank ID: NC_045512.2 and GISAID ID: EPI_ISL_402125) as the outgroup for tree inference. We aligned entire genome sequences by using the FFT-NS-1 program in MAFFT suite v7.407 (Katoh and Standley, 2013) and deleted gapped regions in the 5’ and 3’ regions. We constructed a phylogenetic tree using IQ-TREE 2 v2.1.3 software (Minh et al., 2020) with 1,000 bootstraps (**Figure S1A**). GTR+G substitution model is utilized based on BIC criterion. We found that several sequences without the RSYLTPGD246-253N mutation also clustered with the genomes carrying the RSYLTPGD246-253N mutation (**Figure S1A**), which could be due to reversible mutation(s) and/or recombination (Jackson et al., 2021). Thus, these sequences were excluded from the further analysis.

To estimate the emerging time of the Lambda variant (C.37 lineage), we collected all Lambda sequences carrying the RSYLTPGD246-253N mutation that were sampled in 2020 (2 sequences) and randomly sampled 100 sequences in 2021. We also added the following 4 SARS-CoV-2 genomes as the outgroup: strain Wuhan-Hu-1 (GISAID ID: EPI_ISL_1532199, isolated on December 26, 2019), EPI_ISL_1093172 (isolated on August 25, 2020), and EPI_ISL_1534645 (isolated on November 30, 2020). Note that the two viral genomes isolated in Peru on August 25, 2020 (EPI_ISL_1532199 and EPI_ISL_1093172) were categorized in the B.1.1.1 lineage, although they were previously categorized as the C.37 lineage. We carefully examined these two sequences and found that they could be used as a sister group of the C.37 lineage. As for EPI_ISL_1534645, it does not contain any typical mutations in the S protein, but it the closed to the genuine Lambda variant (Figure S1A). Therefore, we included these four sequences in the analysis. We conducted the Bayesian tip-dating analysis using BEAST v1.10.4 (Suchard et al., 2018). We used GTR+Gamma model for nucleotide substitution model. For the assumption of rate variations, we applied uncorrelated relaxed clock, assuming that the distribution of rates followed a Gamma distribution. We carefully checked the effective sample size of each parameter and confirmed that all are > 200. The emerging time is estimated as 2020.7585 (95% CI, 2020.245–2020.8525). A timetree was summarized using TreeAnnotator software in the BEAST package and visualized by using FigTree v1.4.4 (**Figure 1C** and **Figure S1B**). Reconstruction of the population history, namely the changing on effective population size across time (**Figure 1D**), was conducted by Bayesian skyline plot using the same software and parameter settings using the sampled Lambda sequences as noted in the tip-dating analysis.

#### Protein Structure Homology Model

All protein structural analyses were performed using Discovery Studio 2021 (Dassault Systèmes BIOVIA). In **Figures 2C and 2D**, the crystal structure of SARS-CoV-2 S (PDB: 6ZGE) (Wrobel et al., 2020) was used as the template, and 40 homology models of the SARS-CoV-2 S of the Lambda variant were generated using Build Homology Model protocol MODELLER v9.24 (Fiser et al., 2000). Evaluation of the homology models were performed using PDF total scores and DOPE scores and the best model for the Lambda S was selected. In **Figure S2D**, the cocrystal structure of SARS-CoV-2 and human ACE2 (PDB: 6M17) (Yan et al., 2020) was used.

#### Plasmid Construction

Plasmids expressing the SARS-CoV-2 S proteins of the parental D614G (B.1) (Ozono et al., 2021) and the Epsilon (B.1.427) variant (Motozono et al., 2021b) were prepared in our previous studies. Plasmids expressing the S proteins of the Alpha (B.1.1.7), Beta (B.1.351), Gamma (P.1), Delta (B.1.617.2), Lambda (C.37) variants and the point mutants were generated by site-directed overlap extension PCR using pC-SARS2-S D614G (Ozono et al., 2021) as the template and the following primers listed in **Table S4**. The resulting PCR fragment was digested with Acc65I or KpnI and NotI and inserted into the corresponding site of the pCAGGS vector (Niwa et al., 1991). Nucleotide sequences were determined by DNA sequencing services (Fasmac or Eurofins), and the sequence data were analyzed by Sequencher v5.1 software (Gene Codes Corporation).

#### Pseudovirus Assay

Pseudovirus assay was performed as previously described (Motozono et al., 2021a; Ozono et al., 2021). Briefly, the pseudoviruses, lentivirus (HIV-1)-based, luciferase-expressing reporter viruses pseudotyped with the SARS-CoV-2 S protein and its derivatives, HEK293T cells (1 × 10^6^ cells) were cotransfected with 1 μg of psPAX2-IN/HiBiT (Ozono et al., 2020), 1 μg of pWPI-Luc2 (Ozono et al., 2020), and 500 ng of plasmids expressing parental S or its derivatives using Lipofectamine 3000 (Thermo Fisher Scientific, Cat# L3000015) or PEI Max (Polysciences, Cat# 24765-1) according to the manufacturer’s protocol. At two days posttransfection, the culture supernatants were harvested, centrifuged. The amount of pseudoviruses prepared was quantified using the HiBiT assay as previously described (Ozono et al., 2021; Ozono et al., 2020). The pseudoviruses prepared were stored at –80°C until use. For the experiment, HOS-ACE2 cells and HOS-ACE2/TMPRSS2 cells (10,000 cells/50 μl) were seeded in 96-well plates and infected with 100 μl of the pseudoviruses prepared at 4 different doses. At two days postinfection, the infected cells were lysed with a One-Glo luciferase assay system (Promega, Cat# E6130), and the luminescent signal was measured using a CentroXS3 plate reader (Berthhold Technologies) or GloMax explorer multimode microplate reader 3500 (Promega).

#### Antibody Treatment

Antibody treatment for neutralization and infectivity enhancement were performed as previously described (Saito et al., 2021). Briefly, this assay was performed on HOS-ACE2/TMPRSS2 cells using the SARS-CoV-2 S pseudoviruses expressing luciferase (see “Pseudovirus Assay” above). The SARS-CoV-2 S pseudoviruses (counting ~20,000 relative light unit) were incubated with serially diluted heat-inactivated human sera, a NTD-targeting NAb clone 4A8 (Chi et al., 2020) or an EAb clone COV2-2490 (Liu et al., 2021c)] at 37°C for 1 h. The pseudoviruses without sera/antibodies were included as controls. Then, the 80 μl mixture of pseudovirus and sera/antibodies was added into HOS-ACE2/TMPRSS2 cells (10,000 cells/50 μl) in a 96-well white plate and the luminescence was measured as described above (see “Pseudovirus Assay” above). 50% neutralization titer was calculated using Prism 9 (GraphPad Software).

### QUANTIFICATION AND STATISTICAL ANALYSIS

Data analyses were performed using Prism 9 (GraphPad Software). Data are presented as average with SD.

